# Modelling the Dynamics of Senescence Spread

**DOI:** 10.1101/2023.03.16.532939

**Authors:** Lucy Martin, Linus Schumacher, Tamir Chandra

**Affiliations:** Institute of Genetics and Caner, University of Edinburgh; Centre for Regenerative Medicine, University of Edinburgh

**Keywords:** SASP, Senescence, Mathematical model, Cell signalling

## Abstract

Cellular senescence is a cell surveillance mechanism that arrests the cell cycle in damaged cells. The senescent phenotype can spread from cell to cell through paracrine and juxtacrine signalling, but the dynamics of this process are not well understood. Although senescent cells are important in ageing, wound healing, and cancer, it is unclear how the spread of senescence is contained in senescent lesions. In the absence of the immune system, senescence could theoretically spread infinitely from one cell to another, but this contradicts experimental evidence. To investigate this issue, we developed both a minimal mathematical model and a stochastic simulation of senescence spread. Our results suggest that differences in the number of signalling molecules secreted between subtypes of senescent cells can limit the spread of senescence. We found that dynamic, time-dependent paracrine signalling prevents the uncontrolled spread of senescence and we demonstrate how model parameters can be determined using Bayesian inference in a proposed experiment.

## 1 Introduction

Senescence is a cellular stress response that can be activated by telomere shortening, DNA damage, and the up-regulation of oncogenes (d’Adda di Fagagna et al., 2003; Hayflick & Moorhead, 1961; Serrano et al., 1997). Senescent cells display cell cycle arrest and several changes to the cell, including large-scale chromatin rearrangement and the production of secretory proteins which can induce senescence in nearby cells (the SASP, senescence associate secretory phenotype)(Acosta et al., 2008; Basisty et al., 2020; Coppé et al., 2008; Kuilman et al., 2008; Narita et al., 2003). Senescence is necessary for many processes in tissues including development, wound healing, and defence against tumour formation (Demaria et al., 2014; Kirschner et al., 2020; Muñoz-Espín & Serrano, 2014; Passos et al., 2009). When senescent cells accumulate, the molecules they release can create a pro-tumourigenic and inflammatory microenvironment, which has been linked to both the ageing process and the recurrence of cancerous tumours following treatment (Wang et al., 2020).

SASP molecules can spread the senescence phenotype through paracrine signalling, with molecules diffusing from a secreting senescent cell and binding to non-senescent cells. Several experiments have demonstrated paracrine transmission of senescence, for example, senescent cells contained within a transwell induce senescence in a population of growing cells sharing the same media (Acosta et al., 2013). SASP signalling also recruits immune cells which can clear senescent cells (Prata et al., 2018).

The terms ‘primary’ and ‘secondary’ senescence are used to distinguish between cells in which senescence has been induced cell-intrinsically (e.g, through DNA damage) and extrinsically (through signals from primary senescent cells), respectively (Fig. 1b). Recent evidence shows that these subtypes of senescence are transcriptomically distinct (Rattanavirotkul et al., 2020; Teo et al., 2019).

**Figure 1:**
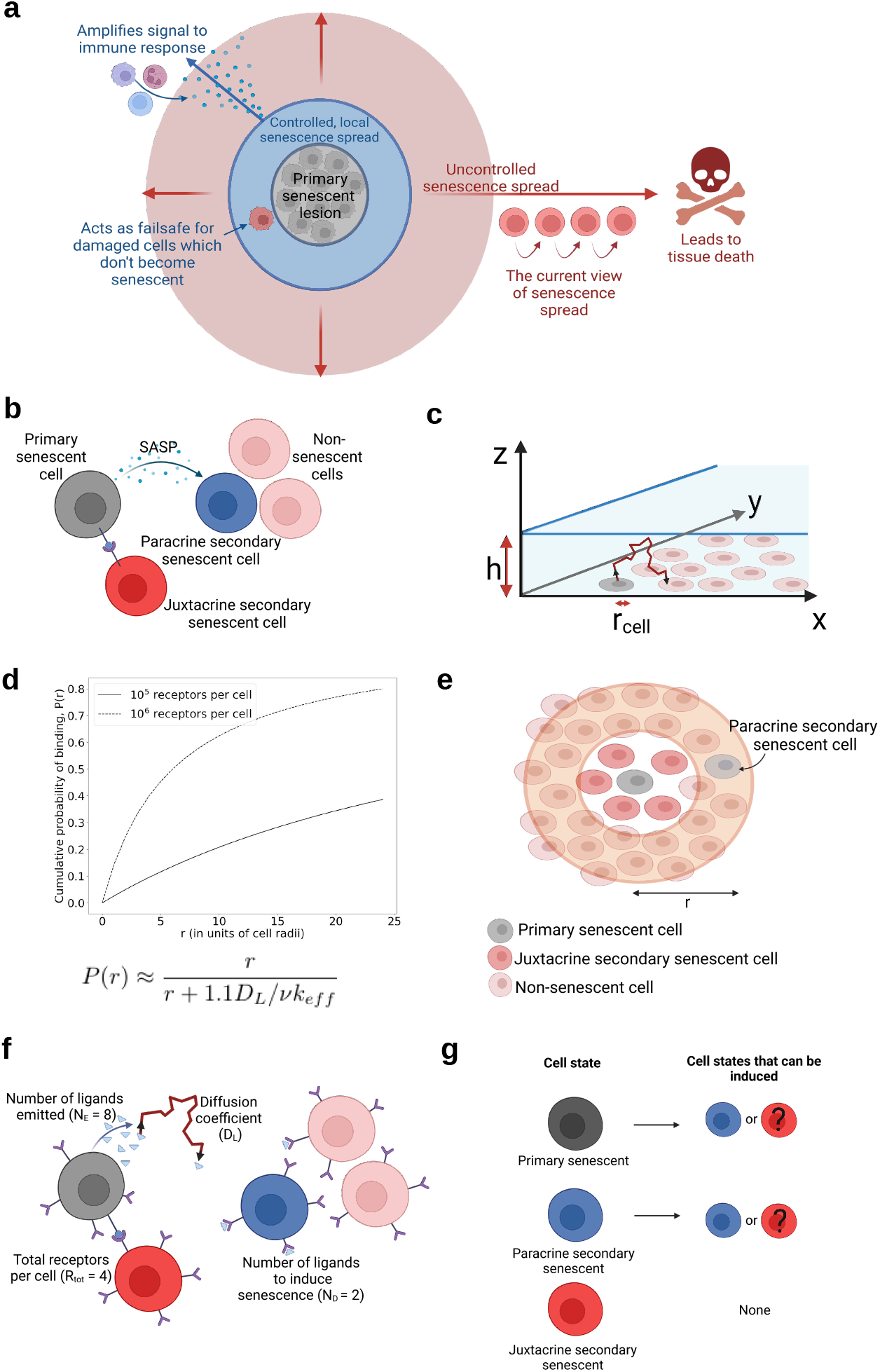
Modeling the spread of senescence. a) A controlled senescence spread is beneficial *in vivo*, but it is unclear how a limited spread of senescence is achieved. b) Illustration of the cellular phenotypes considered in our model. c) A cartoon of the simulated diffusion of ligands through a 3D layer of media over a 2D layer of cells. d) The cumulative probability of a ligand binding at a distance, *r*, from the cell that emitted it. e) An illustration of the minimal model of senescence spread developed in this paper, where we consider the probability of inducing senescence in the orange annulus. f) An illustration of selected model parameters. g) The ways senescent cells can induce senescence in the cells around them.

In addition to paracrine signalling (SASP), senescence can also spread via juxtacrine communication between cells which are in direct contact (NOTCH-mediated) (Hoare et al., 2016; Teo et al., 2019). Secondary senescent cells created through juxtacrine signalling may differ from secondary senescent cells created through paracrine signalling, with juxtacrine secondary cells producing fewer SASP proteins, alongside other transcriptional differences (Hoare et al., 2016; Rattanavirotkul et al., 2020). Whether these two types of secondary senescent cells play different roles in the spread of senescence is as yet unclear (Kirschner et al., 2020).

*In vivo,* a limited and controlled senescence spread is thought to play an important role in at least two scenarios. Firstly, any secondary senescent cells created around a primary senescent lesion may help to amplify the signal to the immune system. Secondly, as a failsafe mechanism, damaged cells that failed to undergo primary senescence will become senescent in response to the primary cells (Fig. 1a). However, multiple lines of evidence suggest that an accumulation of senescent cells and uncontrolled spread will ultimately lead to tissue dysfunction (Baker et al., 2016; Baker et al., 2011; Burd et al., 2013; Katzir et al., 2021).

Currently, there are no hypothesised mechanisms to contain the spread of senescence in the absence of the immune system. However, it has been shown that the spread of senescence is contained when immune cells are not present (Acosta et al., 2013). *In vitro* experiments have shown that from a seeded circle of senescent cells, the spread of senescence is finite (senescence was found at a distance of approximately 1mm from seeded cells) (Acosta et al., 2013). Although medium taken from primary senescent fibroblasts is capable of inducing secondary senescence in growing fibroblasts, it is uncertain if tertiary senescence is induced when the process is repeated (Acosta et al., 2013). Controlled senescence spread is also observed *in vivo*, where some senescent lesions persist over time (melanocytic naevi), and are not cleared by the immune system, but the senescence phenotype does not continue to spread (Bartkova et al., 2006). This is not due to SASP production halting, as cells continue to produce a SASP even months after the onset of senescence and enter a deep senescence state, which can lead to chronic senescence *in vivo* (van Deursen, 2014).

In this paper, we use both a minimal mathematical model and stochastic simulation to investigate the mechanisms that determine the dynamics of senescence spread. We identify that juxtacrine-induced senescent cells can limit senescence spread by acting as a fire-break in the wildfire of paracrine signalling and that this effect is more pronounced when SASP-signalling is delayed compared to juxtacrine signalling. We propose an *in vitro* experiment and demonstrate that the parameters of senescence spread can be inferred using Bayesian statistics. With the emergence of large-scale descriptive efforts such as the SenNet consortium (US) (Lee et al., 2022) and the advancement of spatial transcriptomics (Kiss et al., 2022), our model can serve as a framework for further exploration as new data becomes available.

## 2 Results

### 2.1 Senescence spread model

We simulated senescence spread in a 2D layer of cells covered by a 3D layer of media so that we could compare our results to senescence spread *in vitro*, as senescence is best characterised in adherent human fibroblasts (Cristofalo et al., 2003; Hayflick & Moorhead, 1961).

To simplify the system, we described the cells as circular and non-motile, evenly and randomly distributed on the 2D plane with a specified density. Following work by Batsilas et al.(Batsilas et al., 2003), we considered the cells to be covered by a layer of media of height *h* (as shown in figure 1c). The density of the cells in the 2D plane is given by *σ*, and the radius of each cell is given by *r*_cell_. The number of receptors per cell is *R*_tot_.

Paracrine signalling is mediated by ligands, and in our model the ligands are SASP molecules. Ligands are emitted by senescent cells and diffuse through the surrounding media. *D*_L_ is the diffusion coefficient for the ligand, and a larger diffusion coefficient results in ligands travelling further in a given time frame. Three rates are used to describe a ligand binding to a receptor; *κ*_on_ gives the rate at which ligands bind to a receptor, *κ*_e_ the rate of endocytosis of bound ligand-receptor complexes, and *κ*_off_ the rate at which ligands dissociate from receptors.

Batsilas et al. describe the cumulative probability that a ligand binds at a distance, *r*, from the cell that emitted it, as

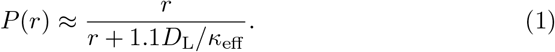

Here, *κ*_eff_ is the effective binding rate of ligands to cells (defined in table 2). As *κ*_eff_ depends on several parameters, including the number of receptors per cell, the cumulative binding probability changes with *R*_tot_ and other system parameters (Fig. 1d).

Building on this work, we assume that cells have a fixed number of receptors for detecting ligands, senescent cells secrete ligands at a fixed rate (*N*_E_), and that a cell must bind a certain number of ligands (*N*_D_) to become senescent. We calculate the rate that ligands bind to a cell at distance *r* from the emitting cell, and the rate that senescence is induced in cells at distance *r* from a senescent cell.

Throughout this paper, we assume that juxtacrine secondary senescent cells do not produce a SASP and cannot in turn induce juxtacrine senescence in their neighbours as a blunted SASP response in juxtacrine secondary cells has been reported (Hoare et al., 2016; Teo et al., 2019). We explore different scenarios of senescence spread, such as secondary or tertiary induction of juxtacrine senescence from paracrine senescent cells, as summarised in figure 1g. We assume that juxtacrine-mediated senescence induction occurs when cells are in contact, i.e., when they are in close spatial proximity.

In a minimal model, we calculate the probability of senescence spread from a single senescent cell, by considering the probability of inducing senescence in an annulus of non-senescent cells around the primary senescent cell, as shown in figure 1e. By considering the probability of senescence spread to a series of annuli, each with the same width, but an increasing radius, we determine the likelihood that senescence spreads from a single cell.

To model more realistic experimental scenarios we built a stochastic simulation of senescence spreading from a collection of cells over time. We use this to investigate the role of secondary senescent cells, by varying parameters such as their SASP secretion, time delays involved in senescence induction, and lesion size.

Detailed descriptions of the minimal model and stochastic simulations can be found in Methods (section 4).

#### 2.1.1 Model parameters

Our model of the spread of senescence contains several parameters, such as the constants relating to ligand-receptor interactions, where exact values have not been determined experimentally. Where this was the case we used values from closely related biological systems, for example, the parameters for EGF and its receptor suggested in Batsilas et al. (Batsilas et al., 2003). These parameters are shown in table 1.

**Table 1:** Table showing the estimated parameters for the spread of EGF, as described in Batsilas et al. (Batsilas et al., 2003), originally from (Wiley et al., 2003).

| Parameter | Meaning | Estimated value |
| --- | --- | --- |
| $N_E$ | Number of ligands emitted per senescent cell per unit time ($\text{h}^{-1}$) | Variable |
| $N_D$ | Number of ligands required to induce senescence per unit time ($\text{h}^{-1}$) | Unknown |
| $D_L$ | Diffusion coefficient for ligands ($\text{cm}^2/\text{s}$) | $10^{-6}$ |
| $\sigma$ | Density of cells ($\text{m}^2/\text{m}^2$) | 0.1 - 0.4 |
| $r_{\text{cell}}$ | Radius of cells (m) | $10 \times 10^{-6}$ |
| $R_{\text{tot}}$ | Total number of receptors per cell | $10^4 - 10^6$ |
| $\kappa_e$ | Endocytosis rate ($\text{min}^{-1}$) | 0.1 - 0.3 |
| $\kappa_{\text{off}}$ | Dissociation rate ($\text{min}^{-1}$) | 0.1 - 0.3 |
| $\kappa_{\text{on}}$ | Forward binding rate constant ($\text{M}^{-1} \text{min}^{-1}$) | $10^8$ |

We assume that a non-senescent cell needs to bind a threshold number of ligands (*N*_D_) in a given time frame for senescence to be induced (Fig. 1f), and that a single senescent cell produces *N*_E_ ligands per unit time. We calculated the number of SASP proteins produced by a senescent cell per hour (Fig. S7) based on a recent paper by Schafer et al. (Schafer et al., 2020) that measured the concentrations of SASP proteins produced by radiation-induced senescent cells. For fibroblasts, the data suggest ~275000 protein molecules per hour across all SASP proteins investigated. However, if we limit the proteins of interest to Interleukin 6 (IL-6), Interleukin 8 (IL-8) and Activin A (secreted proteins thought to be responsible for the induction of senescence (Acosta et al., 2008; DA & PLJ, 2021; Ito et al., 2017; Jochems et al., 2021; Prata et al., 2018)), the number produced per hour is much smaller, ~2887.

For simplicity, we did not distinguish between the types of ligands produced. Although realistically, a specific ligand will bind to its complementary receptor, and multiple signalling molecules may be required to induce senescence, we did not include this complexity in the model.

### 2.2 Cell communication distances are determined by the number of receptors per cell and the effective binding rate

To understand the impact of each model parameter on senescence spread we used our minimal model (described in section 4.1) to scan through a range of parameters, noting the probability of senescence spread from a single primary senescent cell. We chose to fix the number of ligands emitted per hour by a cell (justification in section 2.1.1), and vary *N*_D_, the number of ligands required to induce senescence. For small *N*_D_ we observe uncontrolled spread (blue region in Fig. 2a) whereas high *N*_D_ invariably prevents spread (red region in Fig. 2a). Parameter regimes for a controlled spread are likely to be found at the transition between these regions. We therefore investigated how changing other parameters shifts the values of *N*_D_ for which senescence spreads. Parameter values which increase the maximum of *N*_D_ under which senescence can be induced in a nearby cell correspond to senescence being more likely to spread through the tissue (Fig. 2a). Using this method we were able to draw several conclusions about the dynamics of paracrine signalling in a 2D layer of cells.

**Figure 2:**
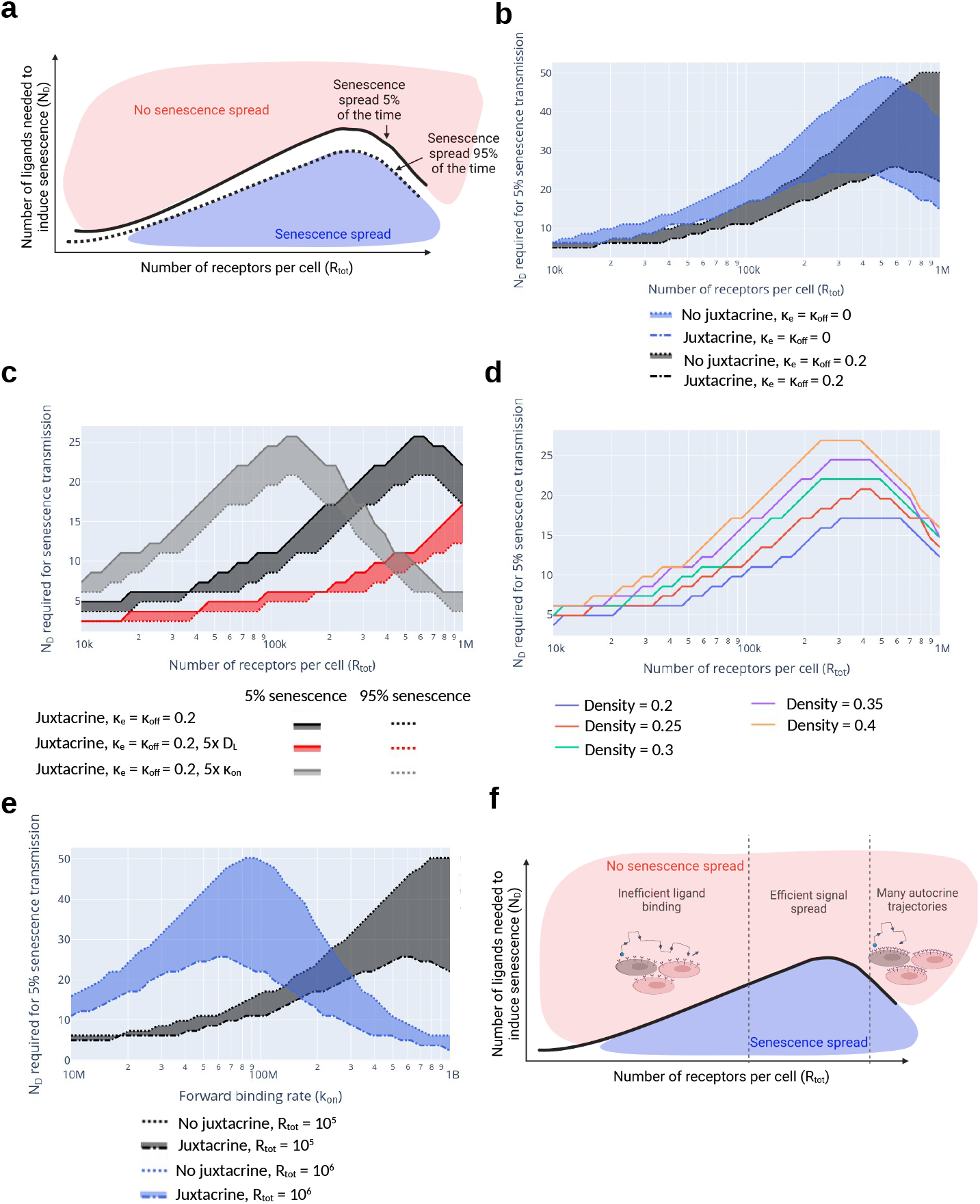
A minimal model for senescence spread. a) An illustration of the phase space of *R*_tot_ and *N*_D_, including a region where senescence spreads and one where senescence does not. b) The division in phase space between senescence spread and no senescence spread for two values of *κ*_e_ and *κ*_off_, with and without juxtacrine senescence. The shaded region shows the parameter regime where senescence would spread without juxtacrine senescence, but not with juxtacrine senescence. c) The range of parameters (shaded) for which senescence only spreads a fraction of the time. d) The division in phase space between senescence spread and no senescence spread for a range of cell densities. e) The division in phase space between senescence spread and no senescence spread for two values of *R*_tot_, with and without juxtacrine senescence. The shaded region shows the parameter regime where senescence would spread without juxtacrine senescence, but not with juxtacrine senescence. f) An illustration of the impact of receptor number on senescence spread. In this figure, unless specified, parameters were, *D*_L_ = 10^-6^ cm^2^/s, *r*_cell_ = 10*μ*m, *κ*_on_ = 10^8^, *σ* = 0.35.

We found that there is both an optimum number of receptors (*R*_tot_) and an optimum forward binding rate (*κ*_on_) for the spread of a signal (Fig. 2(a - e)) in the regime where both *κ*_e_ and *κ*_off_ (the endocytosis and dissociation rates) are negligible, such that ligands which bind to a receptor remain bound. Intuitively, it may seem that more receptors would increase the signal spread, but this is not the case. When ligands do not dissociate from a cell easily, once they are bound they are removed from the system. Therefore, in regimes with large numbers of receptors, ligands are more likely to bind to the cell that emitted them (known as autocrine signalling) or to bind in spatial proximity, so the signal does not spread. Conversely, in regimes with a small number of receptors, the signal diffuses to a much greater distance from the senescent cell, but it is unlikely that enough ligands will bind to a cell to signal effectively (Shvartsman et al., 2001). This is illustrated in figure 2f.

The same logic applies if *κ*_e_ >> *κ*_off_ such that the probability or internalisation, *v*, tends to 1, meaning that the rate of dissociation is relatively low. If the binding rate is large, many ligands will be sequestered in autocrine trajectories, spreading the signal ineffectively. However, if the binding rate is small it is unlikely that enough ligands will bind to a cell to induce a change. Figure 2e illustrates the effect of increasing the binding rate.

When ligands can dissociate effectively and quickly from cells, the optimal number of receptors (for the signal to spread) is increased (Fig. 2b; black vs blue curves). In this regime, a single ligand can bind to multiple cells over the course of its lifetime. Thus we assume only the last binding event is relevant for senescence induction. In this regime, senescence spreads further with a larger number of receptors and a greater binding rate.

A similar effect is observed when increasing the diffusion coefficient (Fig. 2c). With increased diffusivity a ligand can more easily move away from the cell that emitted it, there are fewer autocrine binding events, and so a larger number of receptors per cell can be accommodated without limiting the signal spread.

Compared to the binding rate and receptor number, the cell density has only a small impact on the system’s dynamics, with a greater cell density resulting in a greater chance of senescence spread (Fig. 2d).

### 2.3 Juxtacrine senescence cells limit the spread of senescence

To investigate the effect of juxtacrine senescent cells on the spread of senescence, we first considered a ring of juxtacrine cells around the primary senescent cell in our minimal model. We assume juxtacrine secondary cells do not emit a SASP (Fig. 1g). As expected, the layer of juxtacrine cells reduces the spread of senescence (Fig. 2b; solid vs dotted curves), as ligands emitted by the primary senescent cell bind to the juxtacrine senescent cells, which are already senescent, but in this approximation do not emit ligands of their own. Thus, there is a parameter regime where senescence spread would occur without juxtacrine cells, but does not when they are present (Fig. 2b, shaded region).

To further investigate the effect of juxtacrine senescent cells on the spread of senescence over time, we created a stochastic simulation (described in section 4.2) that also allowed us to investigate the spread from multiple primary senescent cells. Compared to the spread of senescence from a single primary senescent cell, senescence can spread at higher values of *N*_D_, and the secondary senescent cells are more densely distributed around the lesion (Fig. S1). We found that juxtacrine senescent cells also slow the spread of senescence in this more complex setting (Fig. 3), from hours to several days, for a given value of *N*_D_ (Fig. 3c, solid vs dotted curves). However, once senescence starts to spread, it spreads rapidly, leading to an uncontrolled spread. As juxtacrine senescent cells in our model do not produce a SASP, their presence is analogous to removing trees to prevent the spread of a forest fire.

**Figure 3:**
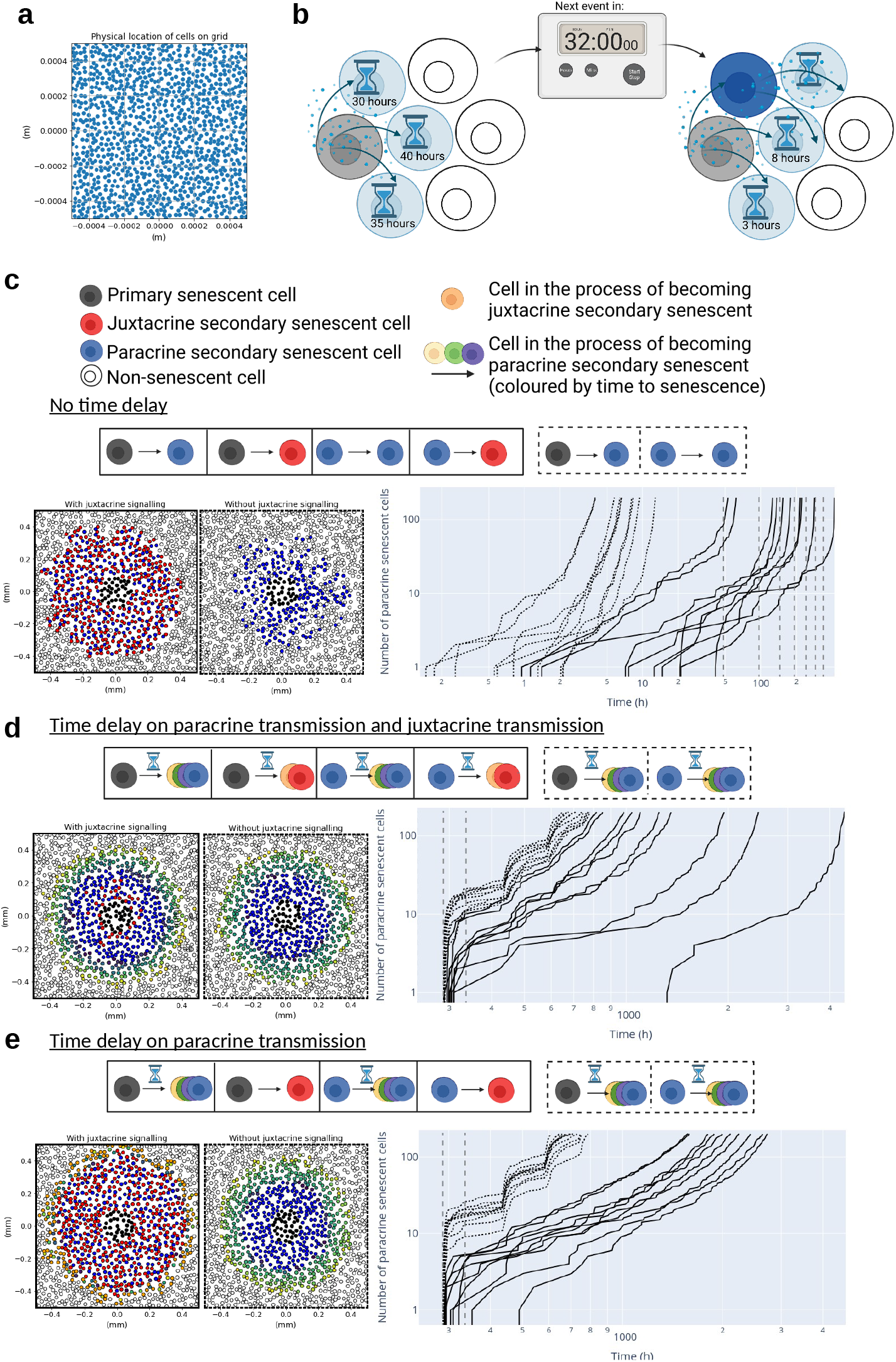
Stochastic simulation of the spread of senescence. a) The simulation grid randomly populated with cells. b) An illustration of the time delay in senescence induction. c) The spread of senescence from a lesion of primary senescent cells with *R*_tot_ = 10^5^, *σ* = 0.35, *k*_e_ = *k*_off_ = 0.2, *N*_D_ = 62, and no delay in the induction of senescence. The left-hand side plots show the spatial distribution of senescent cells and the right-hand side shows the creation of paracrine secondary senescent cells over time, both with juxtacrine senescence (solid) and without (dotted). Vertical grey dashed lines mark days 1-7. d) The same simulation as in (c), but with a 6-day time delay before a cell can induce senescence in its neighbours. e) The same simulation as in (d), but with immediate induction of juxtacrine senescence and delayed induction of paracrine senescence.

### 2.4 A time delay in the senescence induction process limits spread and alters dynamics

After a growing cell binds the required number of ligands to induce senescence, it takes a number of days before the cell becomes fully senescent and it is capable of inducing senescence in its neighbours. To investigate this effect, we introduced a 6-day time delay between senescence induction and a cell becoming fully senescent in the stochastic simulation, as Young et al. suggest 6 days are required to establish full Ras-induced senescence (Fig. 3b) (Young et al., 2009).

We found that introducing a time delay drastically increases the time taken for senescence to spread, as expected (Fig. 3d and S1b). An initial region of paracrine secondary senescent cells is created by the primary lesion, limited in diameter by the reach of the SASP from the primary senescent cells (Fig. 3d). Once these cells become fully senescent and produce their own SASP, the signal spreads further, inducing a new wave of paracrine secondary senescent cells.

The time delay also alters the density of juxtacrine secondary senescent cells, reducing their number at larger distances from the primary lesion (Fig. S5(c and d)). This is due to the assumption that a cell must be fully senescent before it can induce either juxtacrine or paracrine senescence in its neighbours. The delay in juxtacrine induction increases the likelihood that neighbouring cells will become paracrine secondary senescent in the intervening time, increasing the influence of the SASP over senescence spread. The reduction in juxtacrine cells further from the senescent lesion means that as senescence spreads further, the rate of senescence spread increases due to the higher ratio of paracrine secondary senescent cells, which is inconsistent with experimental observations in *vitro* and in vivo (Acosta et al., 2013; Michaloglou et al., 2005).

When *N*_D_, the number of ligands that a cell needs to bind to induce senescence, is decreased, the number of paracrine secondary senescent cells plateaus at a larger number for several days, before increasing sharply again once the cells become fully senescent. In an attempt to find a stable system, where the spread of senescence is limited, we increased *N*_D_. Instead of creating a small, stable number of paracrine senescent cells, this led to no senescence spread on reasonable time scales (several years) from this small primary lesion, suggesting that varying *N*_D_ alone cannot lead to controlled senescence spread.

A dynamic senescence induction may be required to create a stable senescent lesion. Several experiments have determined that senescence induction in a cell is a dynamic process, with evidence that NOTCH signalling (required for juxtacrine induction) is first up-regulated before the cell starts to produce cytokines (for paracrine spread) (Hoare et al., 2016). Preliminary evidence suggests that primary cells induce juxtacrine senescence during this intermediate phase (days 3-5) of senescence activation (Teo et al., 2019). This time-dependent senescence induction may ensure a larger number of juxtacrine secondary senescent cells, limiting the spread of senescence.

### 2.5 A dynamic SASP is required to avoid exponential increases in cell numbers at late times

To further investigate the effects of a dynamic SASP we simulated the limiting case, in which senescent cells can induce juxtacrine senescence in their neighbours as soon as they begin the journey towards becoming fully senescent. In reality, evidence suggests that juxtacrine induction may actually be possible after a 2-day delay (JAG1, the NOTCH ligand responsible for juxtacrine senescence induction, is upregulated even after 1 or 2 days (Hoare et al., 2016)), but simulating the limiting case of immediate juxtacrine senescence induction clearly shows the effect of this change on the dynamics of senescence spread.

The dynamic induction of senescence both slows the spread of senescence and stops the explosion in the number of senescent cells as the lesion grows, which was observed without it (Fig. 3d vs 3e).

### 2.6 Reduced SASP from secondary senescent cells can lead to controlled senescence spread

We wanted to explore parameter regimes in which the spread of senescence is controlled over long periods of time, but this was difficult with such a large number of unknown system parameters.

Once senescence spread from a lesion has begun it can only stop if the balance of system parameters is right, for example, if the flux of SASP from the growing lesion is no longer enough to induce senescence. One way to achieve this is to ensure that the number of juxtacrine secondary senescent cells is large enough to dilute the SASP. This is also achieved if paracrine secondary senescent cells produce fewer SASP molecules than primary senescent cells, as suggested by experimental evidence (Acosta et al., 2013). In our simulations, we find that in this case the primary cells in the initial lesion drive most of the senescence spread (Fig. S2a). Some secondary senescent cells are generated over short time scales, but the spread eventually slows down to the point where it remains controlled, even after 100,000 hours (on the order of a human lifespan). When the lesion size is increased the size of the region of secondary cells will also increase (Fig. S2b).

### 2.7 Parameters of paracrine senescence spread can be determined through inference

To estimate the parameters of the model, we propose a similar experiment to those described in Acosta et al. (Acosta et al., 2013), seeding a circle of primary senescent cells and watching the phenotype spread (section 4.3). By focusing on the first wave of senescence induction, we can forgo computationally intensive simulations. Due to the time delay before a cell becomes fully senescent, there is a period of time when a cell is unable to pass the senescent phenotype on to neighbouring cells through paracrine signalling. If it takes cells 6 days to become fully senescent, we can stop the experiment before 6 days. Using a marker to detect cells that are becoming senescent, we can determine the region to which senescence has spread through paracrine signalling (Fig. 4a).

**Figure 4:**
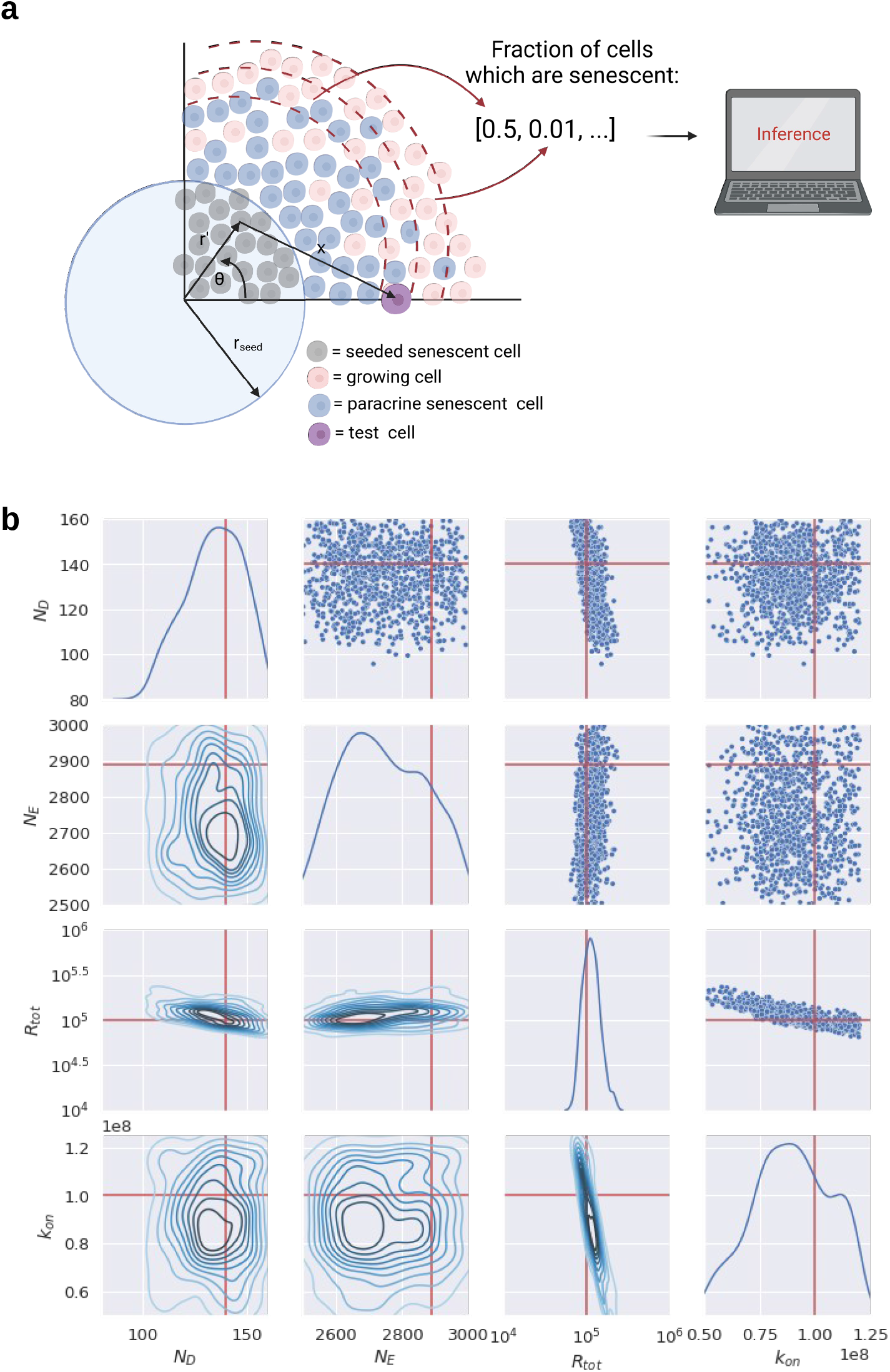
a) A cartoon showing the experimental setup proposed to infer the system parameters. b) Posterior distribution of parameters from the Bayesian inference on simulated data. The plots below the diagonal show the kernel density estimation, and the plots above the diagonal show the final points used in the sampling. Red lines show the true parameter values.

Focusing on short timescales allows us to build on the minimal model described in section 4.1 to predict the fraction of senescent cells at a given distance from the primary lesion. By comparing this with the experimentally observed fraction of senescent cells we can determine *N*_E_, *N*_D_, *R*_tot_, and *k*_on_ using Bayesian inference, fixing other model parameters to experimentally determined values (section 4.3). As a proof of principle for this experiment, we used our model to generate synthetic data and attempted to infer the parameters used to generate the data. As a prior distribution of the parameters, we chose a uniform distribution over the parameter regime given in table 1(Fig. 4b).

We show that it is possible to infer some of the parameters accurately from such an experiment (Fig. 4b). The posterior distribution for *R*_tot_ is narrow and centred on the true value, and although the posterior for *N*_D_ is not as narrow it is well centred on the true value. This reflects the intuition we gained from the minimal model, that senescence spread is sensitive to *N*_D_ and *R*_tot_. The posterior distributions for *N*_E_ and *κ*_on_ are much broader, and less centred on the true values.

## 3 Discussion

Currently, it is unclear how senescence can spread in a controlled way from a primary lesion. We systematically investigated senescence spread, using mathematical modelling informed by experimental evidence. We found that the number of receptors per cell and the binding rate of ligands were critical in determining the distance over which senescence spreads. We observed that an intermediate number of receptors led to the most effective signal spread and uncontrolled senescence spread. Several scenarios (such as juxtacrine-induced cells, in turn, being capable of juxtacrine senescence induction) were not explored as they would lead to the uncontrolled spread of senescence in any parameter regime, highlighting that the current way of thinking about senescence spread is naive.

Our results provide new insights into the role of juxtacrine senescent cells in controlling the spread of senescence. We demonstrate the crucial importance of the dynamic nature of senescence induction in generating juxtacrine secondary senescent cells before paracrine secondary senescent cells and limiting the size of senescent lesions over extended periods. As juxtacrine cells do not emit a SASP (or their SASP is blunted), the presence of juxtacrine senescent cells both reduces the concentration of SASP and limits the parameter regime in which senescence can spread, as their presence limits which cells can become paracrine senescent. Additionally, we propose an experiment aimed at determining four critical parameters involved in senescence spread, and we show that Bayesian inference is a viable tool for estimating these parameters from experimental data.

Although the experiment we propose to quantify the paracrine spread of senescence from a lesion could provide insight into parameters associated with paracrine senescence induction, it will not determine those associated with juxtacrine secondary senescent cells. Conducting such an experiment in practice may be difficult as senescent cells need to be seeded in a circle while controlling the density and movement of surrounding growing cells.

Single-cell transcriptomic data may help to determine the difference between the senescent cell types. However, it is likely that differences between senescent subtypes will vary depending on the cell type and method of senescence induction, suggesting that senescence spread might differ depending on the location in the body and the cause of senescence.

By considering a minimal model of senescence spread we have extracted an understanding of the dynamics of this system. Understanding the way senescence spreads through a population of cells can help us understand several biological processes, from the senescence lesion created in the radiation treatment of cancers to the accumulation of senescent cells as we age. Some questions remain unresolved, such as, whether paracrine senescence cells produce the same amount of SASP as primary senescent cells. These questions may be resolved by especially designed *in vitro* and *in vivo* experiments in combination with our modelling and inference framework.

## 4 Methods

### 4.1 Minimal model

#### 4.1.1 Autocrine and paracrine signalling in cell culture

To understand the limiting factors in the dynamics of senescence spread, we first considered a minimal mathematical model.

To mathematically describe signalling in a 2D layer of cells covered by a 3D layer of media (Fig. 1c), we required knowledge of how a ligand produced by a cell will diffuse through a layer of media. Equation (1), from Batsilas et al. (Batsilas et al., 2003), gives the cumulative probability of a ligand produced by a cell binding at distance *r* from the emitting cell.

To arrive at equation (1), Batsilas et al. assumed that ligands reflect from the upper surface of the media on contact and that ligands can bind to receptors on the surface of cells, but they reflect from the space between cells (the dish). Thus, they approximated the cell-covered dish as a semi-reflecting boundary layer (which is less accurate between 1 < *r/r*_cell_ < 2, close to the emitting cell) and simulated the diffusion and binding of a large number of emitted ligands.

We recreated the simulations described by Batsilas et al. and verified that equation (1) predicts the binding location of ligands with a relative error < 5%. For paracrine trajectories, we compared the time at which a simulated ligand first binds to a new cell to the square of the mean distance at which it binds (Fig. S4b and S4c), showing that the limiting timescale in this process is diffusion (Müller & Schier, 2011). The shape of the cumulative binding probability (*P*(*r*)) is shown in figure 1d, for both small and large numbers of receptors per cell, *R*_tot_.

Equation (1) describes paracrine trajectories, not including ligands that bind to the cell that emitted them (autocrine trajectories). The probability of an autocrine trajectory is given as,

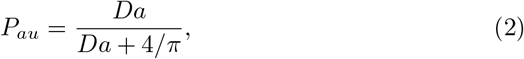

where *Da* is defined in table 2 (Batsilas et al., 2003).

**Table 2:** Table showing compound parameters from (Batsilas et al., 2003).

| Parameter | Meaning | Formula |
| --- | --- | --- |
| $\nu$ | Probability of internalisation | $\kappa_e / (\kappa_e + \kappa_{\text{off}})$ |
| $\kappa$ | Rate constant | $\frac{\kappa_{\text{on}} R_{\text{tot}}}{\pi r_{\text{cell}}^2 N_A}$ |
| $Da$ | Damköhler number | $r_{\text{cell}} \kappa D_L$ |
| $\kappa_{\text{eff}}$ | Effective trapping rate for a homogeneous surface | $\frac{\kappa \sigma}{1 + \pi Da/4}$ |

Equations (1) and (2) do not account for the dissociation of ligands from a cell. If we assume ligands can be internalised by cells via endocytosis with rate *κ*_e_ and dissociate with rate *κ*_off_, the equations can be modified to describe the final distance at which ligands are bound,

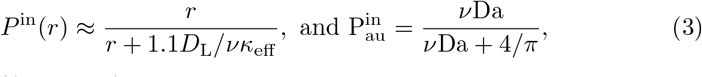

where *v* = *κ*_e_/(*κ*_e_ + *κ*_off_). The first binding event and internalisation are the same if *κ*_off_ = 0, and approach each other when the rate of endocytosis is large and the rate of dissociation small.

Between the first binding event and internalisation a ligand could bind to many cells. However, this binding may be only for a short period of time if the rate of dissociation is high. In this paper we consider only the location of the final binding of the ligand to be important for senescence induction. This assumption breaks down if even fleeting binding of ligands to a cell produces a signal that persists intracellulary, and when combined with the signal from many other fleeting ligand binding events also induces senescence. A comparison of the binding and internalisation distances for an example set of parameters is given in figure S6.

The diffusion of ligands is clearly affected by the height of the media layer, h. However, to sustain cells in *vitro* we would require a layer of media with *h* > 2 mm. Using dimensional analysis and biologically reasonable system parameters, Batsilas et al. (Batsilas et al., 2003) argue that the characteristic trajectory length, *D*_L_/*κ*, is much smaller than 2 mm, and so h does not need to be accounted for.

#### 4.1.2 Senescence spread from a single cell to an annulus

Extending the work of Batsilas et al., we find that for each ligand produced, the probability of a ligand binding in an annulus of width 2*r*_cell_ (illustrated in figure 1e) at distance *r* from the emitting cell is,

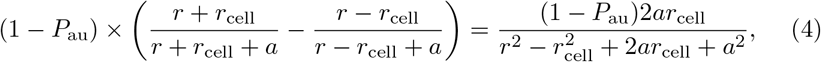

where *a* = 1.1*D*_L_/*νK*_eff_. The area of this annulus is given by 4*πrr*_cell_.

To find the probability that a cell in this annulus will bind the ligand, we multiply this probability by the area of one cell relative to the annulus, to get,

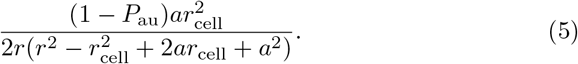

If one cell emits *N*_E_ ligands per unit of time, the expected number of ligands bound to a cell at a distance *r*, per unit of time, is,

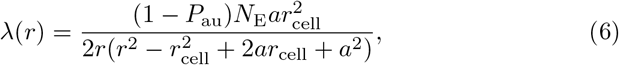

assuming that diffusion has reached a steady state, and *N*_E_ is constant so that the rate of ligands reaching a cell at distance *r* is also constant. This is justified if the diffusion timescale is much faster than the other timescales in the system.

To find the probability per unit time that a specific cell in an annulus will become senescent in any given second, λ_cell_(*r*), we assume that a cell must bind to a threshold number (*N*_D_) of ligands per unit of time in order to become senescent. Here we use the binomial distribution to describe this process, where *N*_E_ becomes the number of trials, and equation (5) gives the probability of success.

However, there is more than one cell in an annulus, on average there will be 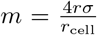. Therefore, λ_spread_(*r*) = *m*λ_cell_(*r*) is the rate of senescence induction in each annulus.

Following this analysis through, we find the probability of creating at least one senescent cell at distance *r* from the primary lesion in time *t* is,

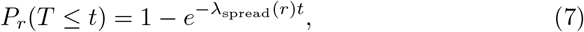

which is 1, minus the probability (given by a Poisson process) that no cells become senescent.

#### 4.1.3 Total senescence spread from a single cell

We now have the probability that one cell at distance *r* from the primary lesion will become senescent in a time period. We use the Poisson-Binomial distribution to calculate the probability that a senescent cell is created in at least one annulus, as the probability of senescence induction in each annulus is different. Therefore, the probability of senescence induction over all annuli is given by a set of independent Bernoulli trials that are not identically distributed, where each annulus is a Bernoulli trial with *p* = *P_r_*(*T* ≤ *t*), and we calculate the probability that one or more of these trials is successful.

We use this simplified model of the system to determine whether the spread of senescence is uncontrollable, considering the limiting case of a single primary senescent cell. If a single senescent cell spreads senescence to one other cell before the body can remove it (we approximate that this happens in 2 days), with a probability of ~1, we define the spread of senescence as uncontrollable. This uncontrollable spread of senescence is not observed *in vitro*, and therefore this is an unrealistic parameter regime.

#### 4.1.4 Senescence spread with juxtacrine cells

We considered two different regimes, with and without juxtacrine secondary senescence. Where juxtacrine signalling induces secondary senescence, we expect that cells surrounding the primary senescent cell will become juxtacrine senescent through contact. Assuming that juxtacrine secondary senescent cells do not emit a SASP, for the senescence to spread, a cell beyond this ring of juxtacrine cells must become senescent within 2 days. If juxtacrine cells instead produce a blunted SASP, this will lead to an underestimate of the spread of senescence (Rattanavirotkul et al., 2020).

The distance of the annulus from the primary senescent cell which is considered in each case is given by,

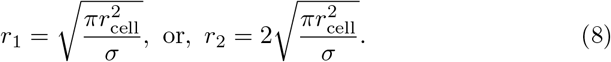

The first value, *r*_1_ is simply the average distance between cells, and *r*_2_ = 2*r*_1_ to account for a ring of juxtacrine senescent cells.

#### 4.1.5 Limitations

This minimal mathematical model provides a tractable way to consider senescence spread from a single cell but is difficult to extend to a system containing many cells. Furthermore, this model cannot be used to investigate the effect of a time delay in senescence induction and it makes a number of assumptions which may limit the accuracy of the predictions.

### 4.2 Stochastic Simulations

In order to deepen our understanding of the system and more closely match the reality of senescence spread *in vivo*, we created a stochastic model for the spread of senescence from multiple initial cells and over many subsequent senescence induction events. The stochastic model lets us introduce the added complexity of paracrine senescent cells which can in turn emit a SASP. We used the simulation to explore the change in dynamics resulting from changing the strength of the SASP produced by these secondary senescent cells. Finally, we introduce a time delay between senescence induction and SASP, more accurately modelling the cellular biology. Cells do not immediately start emitting a SASP once they have bound enough ligands to induce senescence, and instead take several days to become fully senescent.

#### 4.2.1 Simulation implementation

We consider three possible cell states: senescent, in the process of becoming senescent, and non-senescent. A cell starts to become paracrine secondary senescent when it binds to more than *N*_D_ ligands per unit of time (see section 2.1.1). The simulation is initialised on a square plane, and a 100 x 100 cell radii area is commonly used as larger areas increase the computational time taken for each simulation. Cells are then created so that they randomly populate this area with a specified density (Fig. 3a). Primary senescent cells are seeded at time 0. The simulation proceeds by tracking the state of each cell.

In the stochastic simulation, we approximate the number of ligands bound from each emitting senescent cell using the Poisson distribution. This approximation holds when the number of ligands emitted, *N*_E_, is large and the probability of binding (equation (5)) is small. Large binding probabilities are not of interest as they would lead to the fast and uncontrolled spread of senescence. The number of ligands bound from all senescent cells is given by a Poisson distribution whose mean is the sum of the Poisson distribution of ligands from each emitting cell (equation 6),

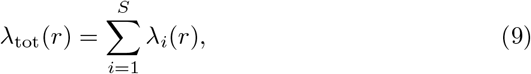

where *S* is the number of senescent cells emitting a SASP.

This approximation dictates the appropriate time frame for the simulation. Considering paracrine signalling using only IL-6, IL-8, and Activin A molecules we can estimate *N*_E_ ≈ 2887 *h*^-1^.

We can now calculate the probability of a cell becoming senescent in one unit of time from its position relative to the SASP-emitting cells. This probability can then be used to give the rate of senescence induction for that cell. Thus, each cell on the grid which is not yet senescent has a rate of senescence induction. By summing these rates we calculate the time at which the next cell becomes paracrine secondary senescent. We use the Gillespie algorithm, meaning that we first determine the time an event occurs based on creation rates, for example, a cell becomes paracrine secondary senescent at t = 1.5h. We then determined which cell becomes senescent, again based on the rate of senescence induction associated with each cell (Gillespie, 1977).

The situation is complicated further by the time delay: It may be that cells have finished becoming fully senescent before the next paracrine induction event, in which case the SASP that they produce must also be accounted for. This is achieved simply by tracking the number of cells that are in the process of becoming senescent. If a cell finishes becoming senescent before the next reaction (i.e., new senescence induction event) occurs, then the cell is added to the list of fully senescent cells, and the simulation run again from that time point, with the probabilities of each cell becoming senescent re-calculated including any SASP from the newest senescent cell. This works as the system is Markovian.

There are a number of algorithms which can be used to speed up the delayed reaction calculation (Cai, 2007). However, at any one time, we do not have a large number of cells waiting to become senescent and the bottleneck in the process is the repeated calculation of the probability of inducing senescence in each cell.

The simulation code can be found on GitHub (https://github.com/lkmartin90/Senescence_Spread).

#### 4.2.2 Comparison of stochastic simulations to minimal model

We compared our stochastic simulation to our minimal model to verify that the two agree. To do this, we considered the spread of senescence from a single cell, in the case where no juxtacrine secondary senescent cells are created.

We used the stochastic simulation to simulate this scenario 100 times (fixing all parameters but initialising a new population of cells each time) and recorded the proportion of the simulations in which there was no senescence spread. We varied *N*_D_ and *R*_tot_, to ensure that the simulations agree over a range of parameters. We compared the results to our minimal model, which gives us the probability of senescence spread.

From this, we see a good agreement between the two methods at both *R*_tot_ = 10^5^ and *R*_tot_ = 10^6^, both when cells in the stochastic simulation are created so that they lie on a lattice (Fig. S3a and S3b), and when cells are randomly distributed (Fig. S3c and S3d). There are some differences in the regime without juxtacrine senescence induction with *R*_tot_ = 10^6^, due to the large number of ligands which bind to nearby cells and the assumption we make when constructing the minimal model that cells are uniformly distributed (Fig. S3b, dotted).

In the stochastic simulation, with cells distributed randomly (Fig. S3c), the range of parameters that result in a probability of senescence spread > 0 and < 1 is greater. If a nearby cell happens to be closer than the average distance between cells then there is a higher probability that senescence will spread. This implies that the parameter regime of large *R*_tot_ is highly dependent on the initialisation of the simulation, in particular, the precise location of the cells. Although the average density in each simulation is the same, the signal spread is possible in some cases and not others. If there are no cells close enough to the primary senescent cell, senescence will not spread.

We investigated whether the optimum number of receptors that we found for senescence spread in the minimal model, as shown in figure 2, also appears in the stochastic simulation, or whether this is a consequence of the simplifying assumptions made in the minimal model. We see that the random nature of the cell placement in the stochastic simulation means that there is no longer a clear optimal number of receptors to ensure the spread of senescence, however, the general dynamics remain the same, with juxtacrine cells reducing the spread of senescence and the number of receptors altering the distance over which cells communicate (Fig. S3d).

#### 4.2.3 Limitations

The stochastic simulation is a useful tool to understand senescence spread from a small lesion, but computationally intensive to scale to larger spatial regions.

### 4.3 Parameter Inference

We use Bayesian statistics to infer system parameters from a realistic *in vitro* experiment, where a circle of primary senescent cells is seeded in the centre of a dish, as described in section 2.7. We calculate the distance senescence spreads from the circular lesion.

The number of SASP molecules emitted per unit area by a primary senescent lesion is, 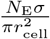, and the probability that a single test cell (Fig. 4a) at distance *x* from a senescent cell binds a ligand emitted by a senescent cell is,

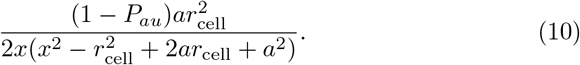

Integrating for the emission from the area of seeded cells to get the total expected number of ligands binding to the test cell gives,

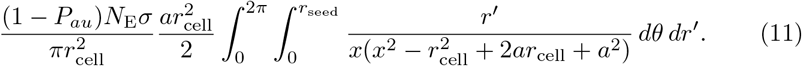

Writing *x* in terms of *r, r*′, and *θ*, according to the geometry in figure 4a, gives,

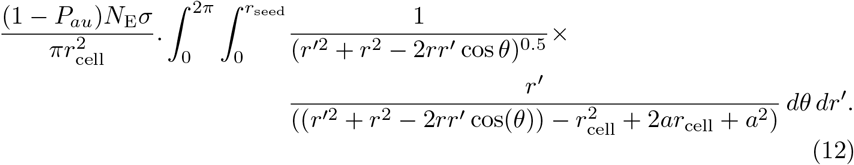

Here *r*_seed_ is the radius of the circle of primary senescent cells. We solve this integral numerically.

Using the method previously described in section 4.1, we calculate the probability that the cell that binds these ligands becomes senescent, using a Poisson distribution to determine the probability that more than *N*_D_ ligands are bound per hour. We then calculate the probability that a cell becomes senescent within the duration of the experiment. Finally, we calculate the fraction of cells which are senescent at a given distance from the seeded senescent region of cells. As this process is probabilistic, the fraction of senescent cells varies on each run of the model.

Before comparing the model output to an experiment, we ought to consider whether the assumptions made in the model’s construction are reasonable. In particular, in the model, we have not accounted for the time taken for ligands to diffuse away from the cell that emitted them. From figure S4, we can see that the overall process of signal spread is diffusion-limited. Previous experiments show senescence spread over a distance of the order of 1 mm, indicating that this is a meaningful distance to consider. The time taken for ligands to travel 1 mm is between 50 and 100 seconds for the parameter regimes shown, depending on the diffusion coefficient of the ligand. As long as the time taken for diffusion is small compared to the timescales associated with senescence spread (hours), we can assume that the diffusion occurs instantaneously.

To determine the parameters which lead to the closest agreement between the model and the experimental data we compare two summary statistics. Firstly, the distance where the fraction of senescent cells crosses 0.5 (*S*). Secondly, the fraction of senescent cells at different distances from the lesion 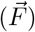. The distances need not be equally spaced, and we use more data points around the region of *r* = *S*, where the fraction of senescent cells is < 1 and > 0. We verified that these summary statistics are sensitive to changes in the parameters (Fig. S8a).

To quantify the agreement between predicted and observed summary statistics, use the distance function:

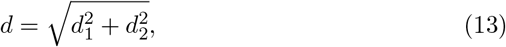

where *d*_1_ = *S*_model_ – *S*_data_ and 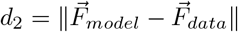.

We use pyABC (Klinger et al., 2018), a likelihood-free inference software using a sequential Monte Carlo scheme, to estimate posterior distributions of model parameters. In this paper, we use synthetic data, created from the same model we use for the inference.

Our inference ran on 60 cores for 13 populations, with 1000 particles, taking 7.5 hours in total. While acceptance thresholds for *d* (epsilon values) have not converged at this stage, indicating that further iterations may improve the inference, the fraction of accepted samples was ≈ 10^-3^, making it computationally wasteful to run for more populations (Fig. S8b).

## Code Availability

Code used in this manuscript is available at https://github.com/lkmartin90/Senescence_Spread

## Contributions

L.S. and T.C. supervised the study. L.K.M., L.S. and T.C. wrote the manuscript. L.K.M. and L.S. developed the modelling framework. L.K.M. conducted the analysis.

## Acknowledgments

L.K.M is a cross-disciplinary research fellow supported by funding from the CRUK Brain Tumour Centre of Excellence Award (C157/A27589). T.C. is supported through a Chancellor’s Fellow at the University of Edinburgh and the MRC Human Genetics Unit. We thank M. Nicholson for critical reading of the manuscript. We thank all members of the Chandra lab and Schumacher lab for their input.

