## Supplementary material for "Modelling the Dynamics of Senescence Spread"

### Supplementary Information

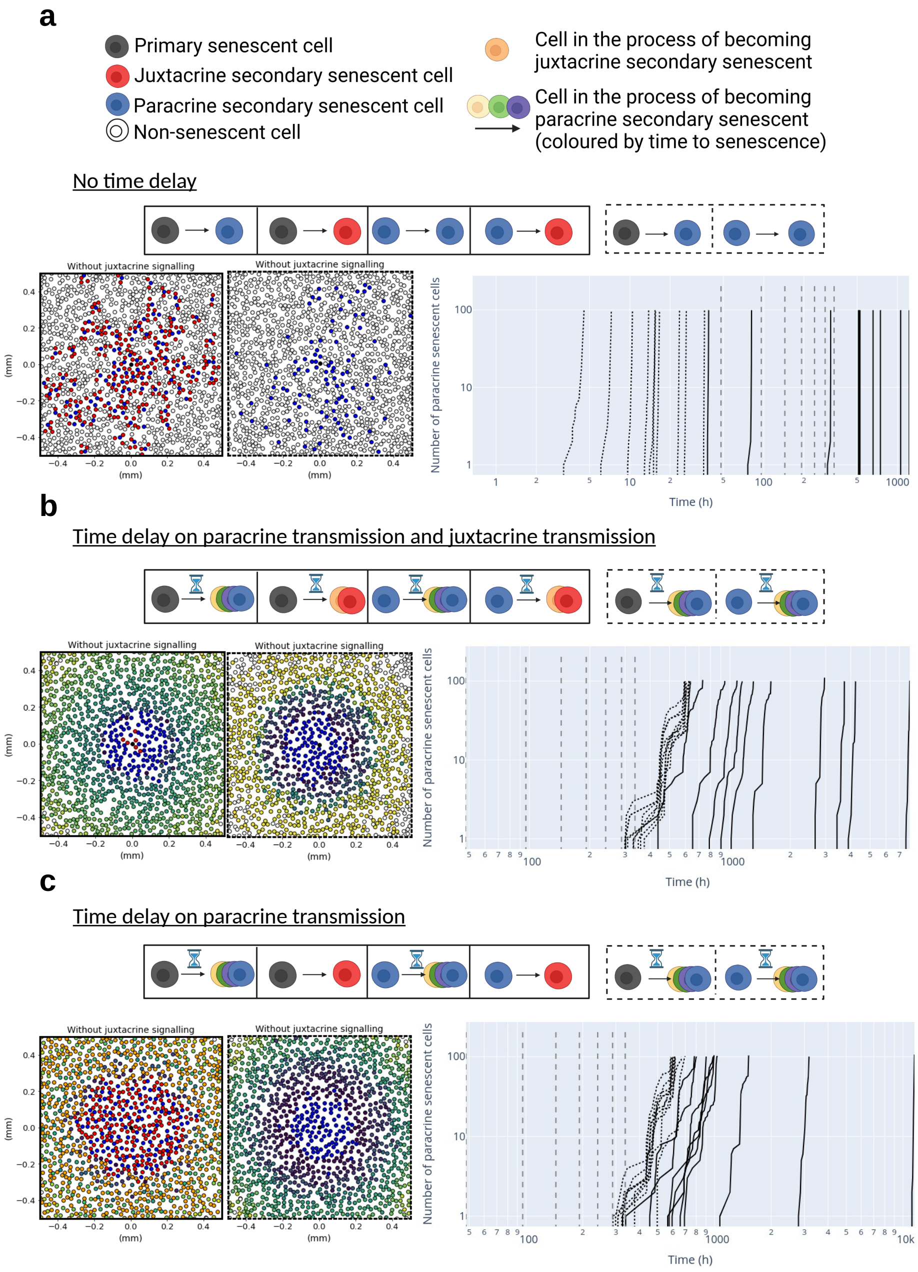


Figure S1:

Figure S1: a) The spread of senescence from a single primary senescent cell with $R_{\mathrm{tot}} = 10^5$, $\sigma = 0.35$, $k_e = k_{off} = 0.2$, $N_{\mathrm{D}} = 9$, and no delay in the induction of senescence. The left-hand side plots show the spatial distribution of senescent cells and the right-hand side shows the creation of paracrine secondary senescent cells over time, both with juxtacrine senescence (solid) and without (dotted). Vertical grey dashed lines mark days 1-7. b) The same simulation as in (a), but with a 6-day time delay before a cell can induce senescence in its neighbours. c) The same simulation as in (b), but with immediate induction of juxtacrine senescence and delayed induction of paracrine senescence.

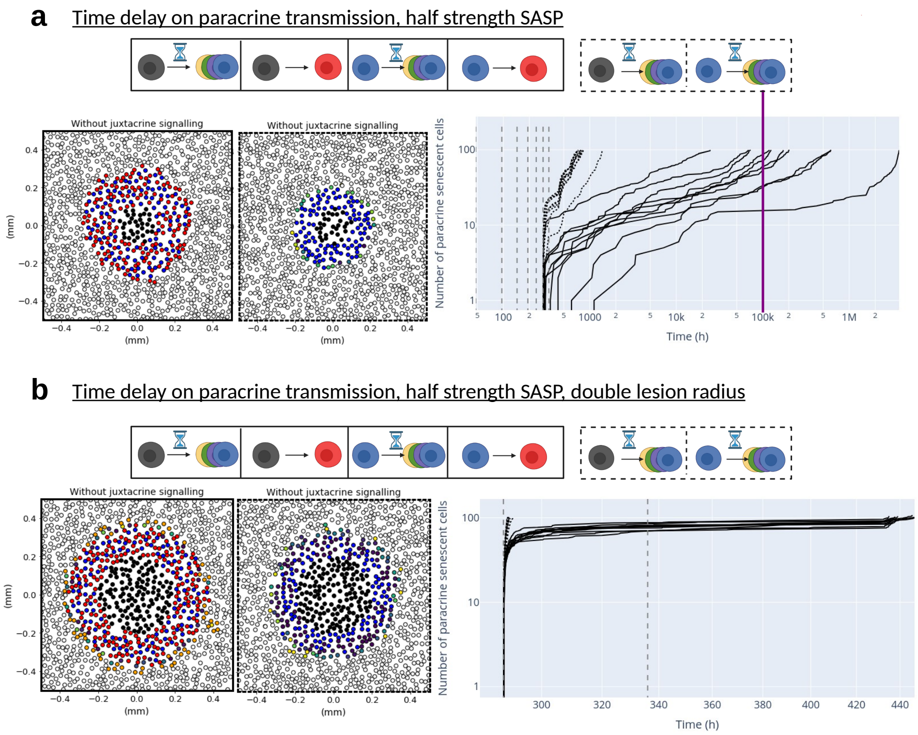


Figure S2: a) The spread of senescence from a lesion of primary senescent cells with $R_{\text{tot}} = 10^5$, $\sigma = 0.35$, $k_e = k_{off} = 0.2$, $N_{\text{D}} = 62$, and a 6-day delay in the induction of senescence. In this simulation, paracrine secondary senescent cells produce half of the SASP of primary senescent cells. The vertical purple line highlights 100,000 hours, which is on the order of a human lifespan. d) The same simulation as in (c), but from a larger primary lesion.

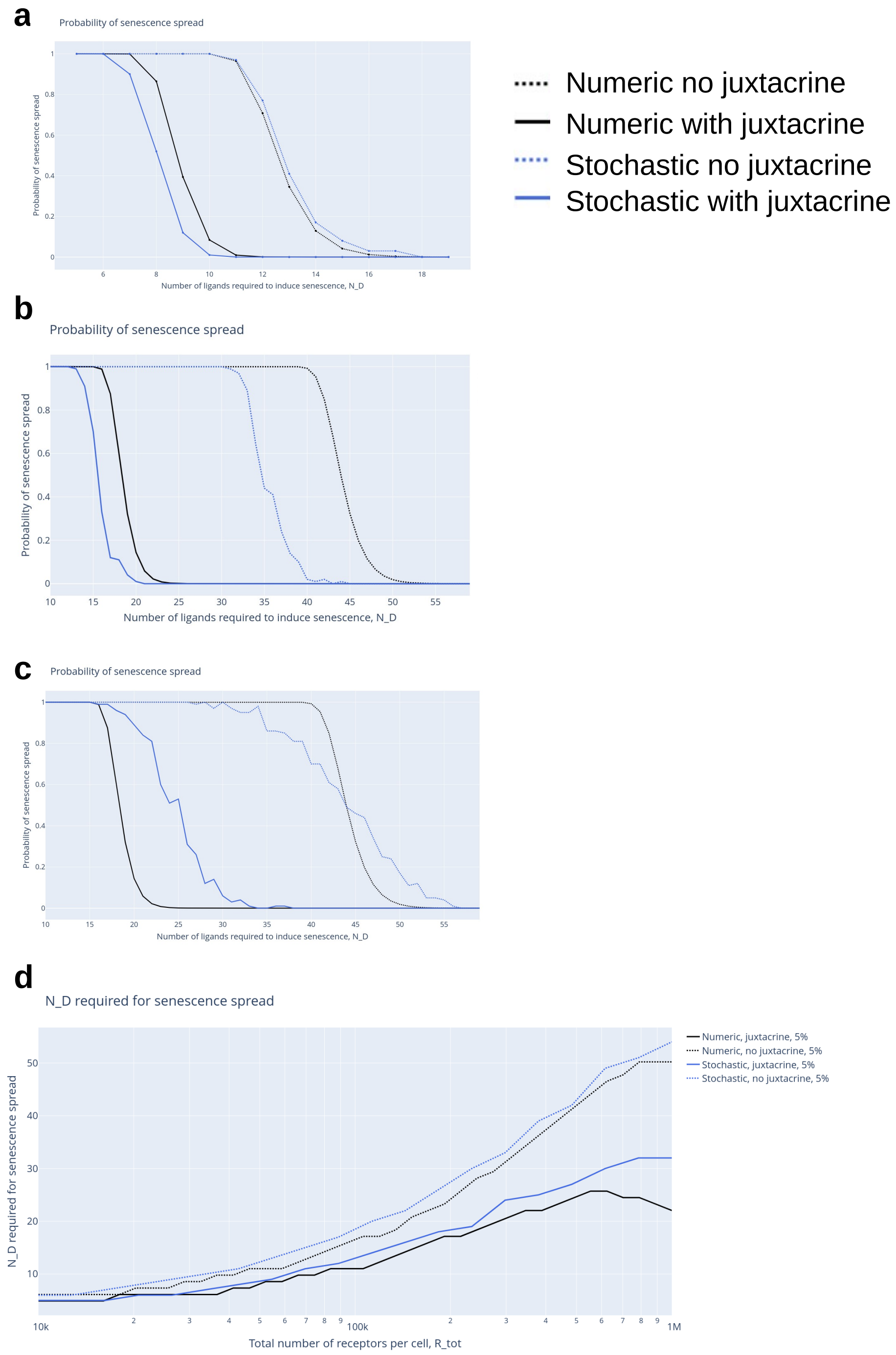
a
Probability of senescence spread
Probability of senescence spread
Number of ligands required to induce senescence, N_D
Numeric no juxtacrine
Numeric with juxtacrine
Stochastic no juxtacrine
Stochastic with juxtacrine
b
Probability of senescence spread
Probability of senescence spread
Number of ligands required to induce senescence, N_D
c
Probability of senescence spread
Probability of senescence spread
Number of ligands required to induce senescence, N_D
d
N_D required for senescence spread
N_D required for senescence spread
Total number of receptors per cell, R_tot
Numeric, juxtacrine, 5%
Numeric, no juxtacrine, 5%
Stochastic, juxtacrine, 5%
Stochastic, no juxtacrine, 5%

Figure S3:

Figure S3: Comparison of the stochastic simulation to the minimal model. In (a) - (d) the stochastic simulation was run 100 times for a single primary senescent cell and the probability of senescence spread in 2 days against the number of ligands required to induce senescence. a) $R_{\text{tot}} = 10^5$, density$= 0.35$, $k_e = k_{off} = 0.2$, $k_{on} = 10^8$, cells are constrained to a lattice. b) A repeat of the simulation in (a) with $R_{\text{tot}} = 10^6$. c) A repeat of (b), but cells are not constrained to a lattice. d) A comparison of the $N_D$ values required to see 5% senescence spread from a single cell in the minimal model and the stochastic simulations.

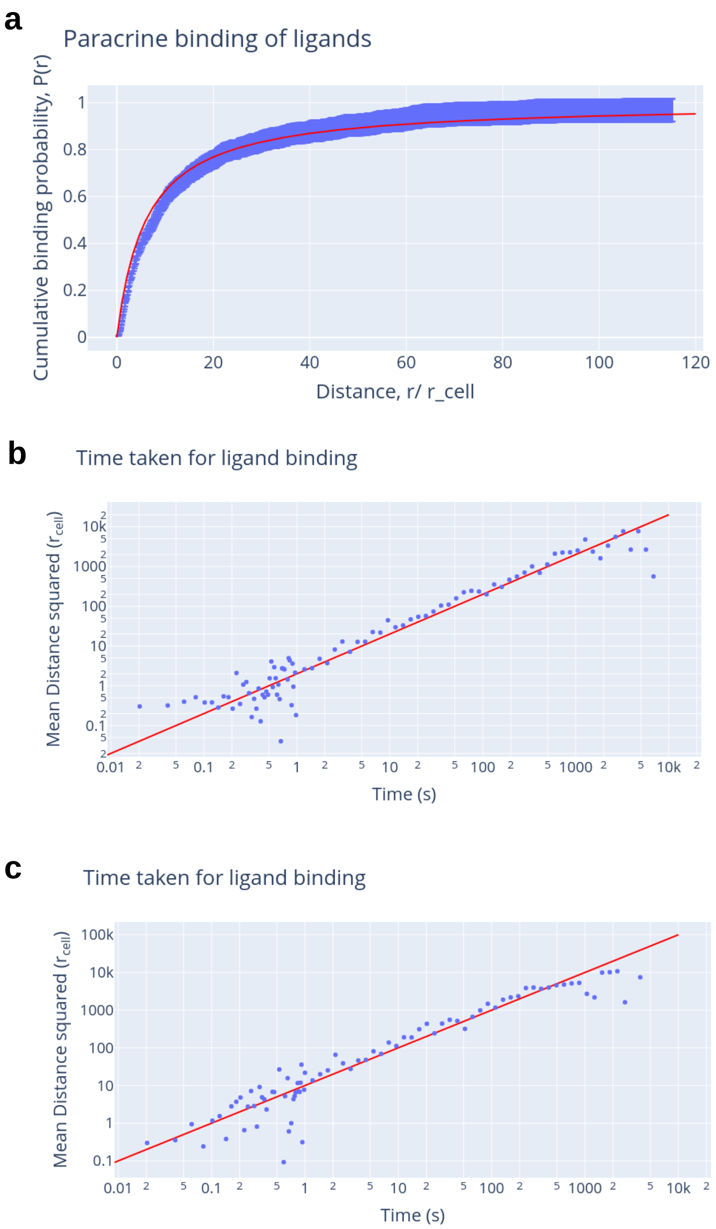
a
Paracrine binding of ligands
Cumulative binding probability, P(r)
Distance, r/ r_cell
b
Time taken for ligand binding
Mean Distance squared (r_cell)
Time (s)
c
Time taken for ligand binding
Mean Distance squared (r_cell)
Time (s)

Figure S4:

Figure S4: Results from the recreation of the ligand binding code described by Batsilas et al, in (Batsilas et al., 2003). 1000 simulations were run in which a ligand was tracked as it was emitted from a cell until the first time it bound to a cell. a) A comparison of the cumulative binding observed in the simulation with a 5% error bar (blue) and equation 1 (red). Both (b) and (c) show the results of these simulations, plotting the mean distance at which a ligand bound against the time it took for this binding to occur. The blue points show the simulation result, and the red line the expected result if this was a diffusion-limited process. a & b) $D_{\mathrm{L}} = 10^{-6}$ cm$^2$/s, cell density = 0.2, media height = 0.002 m, $R_{\mathrm{tot}} = 10^6$, and $k_{on} = 10^8$. b) $D_{\mathrm{L}} = 5 \times 10^{-6}\ cm^2/s$, cell density = 0.35, media height = 0.002 m, $R_{\mathrm{tot}} = 10^5$, and $k_{on} = 10^8$.

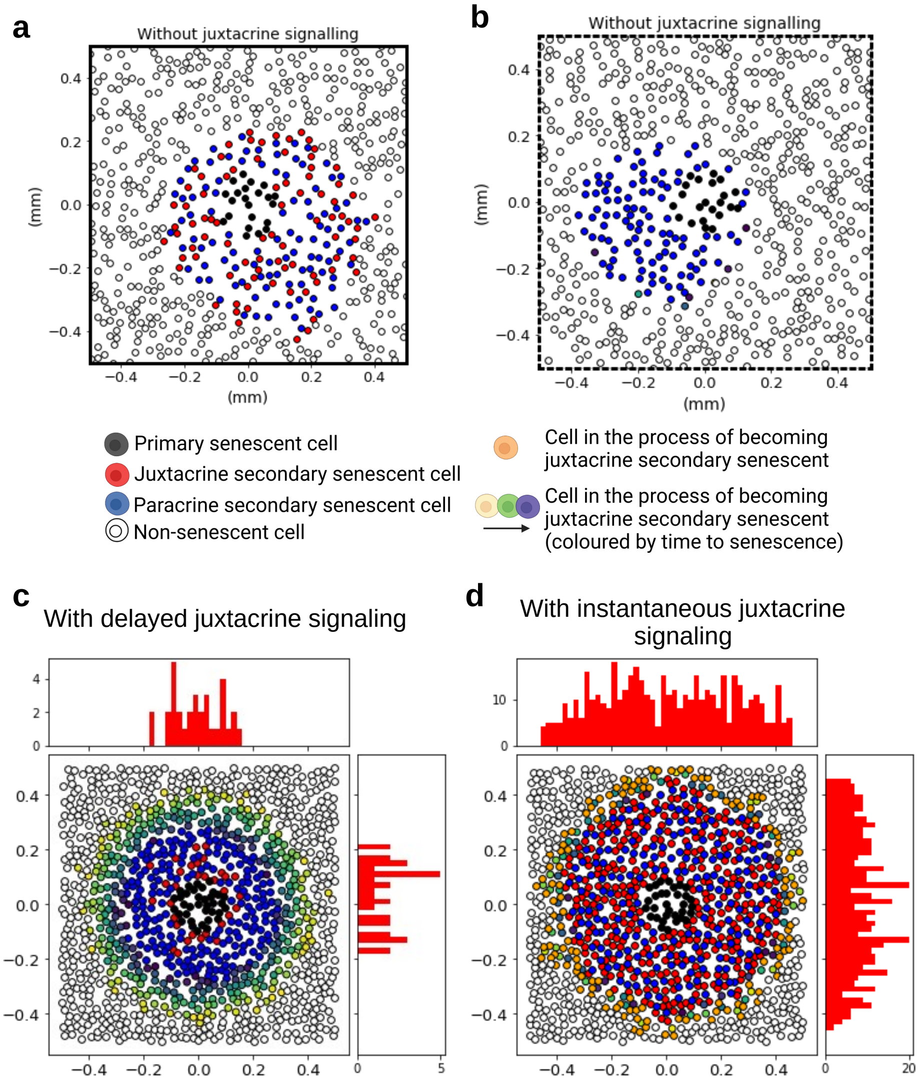


Figure S5: (a) and (b) show a repeat of the simulation in figure 3e, with the reduced density of 0.2. (c) and (d) show senescence spread from a lesion both with and without a delay in the juxtacrine signalling, with histograms showing the density of juxtacrine secondary senescent cells.

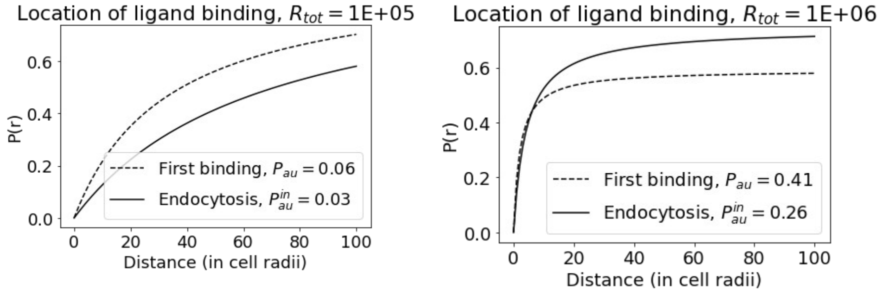


Figure S6: Comparison of the binding and internalisation distances.

| | HUVEC CTRL | HUVEC SnC | Epithelial CTRL | Epithelial SnC | Preadipocytes CTRL | Preadipocytes SnC | Myoblasts CTRL | Myoblasts SnC | Fibroblasts CTRL | Fibroblasts SnC |
|---|---|---|---|---|---|---|---|---|---|---|
| **Activin A** | 3.380553e+02 | 6.032922e+03 | 465.934038 | 1.756599e+03 | 976.211790 | 2.751427e+04 | 214.630247 | 942.863936 | 996.554440 | 1454.315050 |
| **ADAMTS13** | 1.929830e+04 | 1.924970e+05 | 26181.901720 | 1.032388e+05 | 11004.596164 | 1.704250e+05 | 27697.527425 | 25141.001856 | 12715.827592 | 33579.838483 |
| **CCL3** | 3.231604e+03 | 2.329053e+04 | 3738.965306 | 1.967156e+04 | 30794.079733 | 3.866217e+04 | 9547.373944 | 3988.433700 | 19802.804612 | 26283.721014 |
| **CCL4** | 5.495414e+03 | 3.960043e+04 | 7710.569481 | 3.207503e+04 | 14565.719021 | 5.672590e+04 | 12310.214470 | 7873.443957 | 11119.457894 | 24314.415200 |
| **CCL5** | 1.445445e+02 | 1.273918e+03 | 144.301323 | 5.950946e+02 | 109.687882 | 1.219488e+03 | 60.544307 | 155.735313 | 126.642133 | 126.642133 |
| **CCL17** | 3.138710e+03 | 2.542444e+04 | 3936.400516 | 1.634218e+04 | 3658.882759 | 2.716445e+04 | 4701.445582 | 4047.518181 | 3000.088150 | 9486.995670 |
| **CCL22** | 1.105643e+03 | 9.068385e+03 | 1569.185705 | 6.449073e+03 | 3587.119295 | 1.165531e+04 | 2627.508492 | 1614.352703 | 1926.008114 | 5621.400918 |
| **Fas** | 2.111364e+02 | 2.006197e+03 | 480.317716 | 2.035801e+03 | 695.222070 | 2.453039e+03 | 322.303223 | 402.694676 | 249.262128 | 1193.374251 |
| **GDF15** | 1.451260e+03 | 6.089345e+04 | 80.110805 | 4.030566e+02 | 348.195217 | 2.401518e+03 | 210.714692 | 1132.620922 | 200.242188 | 2351.403440 |
| **GDNF** | 1.279061e+01 | 1.321732e+02 | 16.797710 | 6.990011e+01 | 13.818766 | 1.612467e+02 | 22.927376 | 18.870616 | 9.235503 | 37.489335 |
| **ICAM1** | 1.082503e+04 | 8.966639e+04 | 15240.786340 | 6.437089e+04 | 8419.100945 | 1.039911e+05 | 18052.999187 | 14884.019720 | 16868.248433 | 34793.934983 |
| **IL-6** | 2.108345e+02 | 1.406211e+03 | 38.660340 | 1.486155e+02 | 381.388222 | 3.011318e+04 | 4.739725 | 74.151730 | 3.910138 | 220.965626 |
| **IL-7** | 6.898974e+01 | 5.642952e+02 | 94.147628 | 3.907221e+02 | 54.021999 | 6.168430e+02 | 91.052391 | 86.364038 | 13.629760 | 148.776748 |
| **IL-8** | 8.738398e+02 | 4.032366e+04 | 1665.956513 | 6.019060e+03 | 176.355387 | 2.587868e+03 | 101.784804 | 362.272321 | 98.370643 | 1213.629612 |
| **IL-15** | 6.910131e+01 | 5.801723e+02 | 89.079939 | 3.637656e+02 | 3.676000 | 5.144127e+02 | 76.671871 | 74.876639 | 56.042528 | 101.250539 |
| **MMP2** | 2.766824e+04 | 2.853724e+05 | 8831.892628 | 7.033632e+04 | 21770.822708 | 1.195212e+05 | 34140.031777 | 57218.289557 | 8302.101996 | 37834.792949 |
| **MMP9** | 8.838870e+01 | 7.316914e+02 | 121.841588 | 5.029193e+02 | 245.314590 | 7.512010e+02 | 157.817969 | 108.310897 | 182.109921 | 326.647272 |
| **OPN** | 2.375762e+04 | 6.211136e+05 | 29278.518260 | 1.459774e+05 | 48578.947044 | 1.368024e+05 | 27042.968443 | 19545.239759 | 51569.404871 | 71483.807020 |
| **PAI1** | 3.046816e+06 | 8.272197e+07 | 674414.832554 | 2.561970e+06 | 21794.141598 | 1.121929e+06 | 65269.750866 | 156181.784510 | 2228.416155 | 22319.638150 |
| **PAI2** | 9.629501e+01 | 8.602092e+02 | 100.002652 | 3.985709e+02 | 0.157961 | 2.778543e+02 | 5.901568 | 37.115478 | 0.150558 | 0.150558 |
| **SOST** | 3.609486e+01 | 3.423851e+02 | 44.069395 | 1.685227e+02 | 57.213207 | 1.916424e+02 | 14.612258 | 43.375532 | 37.245224 | 37.245224 |
| **TNFa** | 1.697092e+01 | 1.388752e+02 | 21.036692 | 9.085795e+01 | 10.656905 | 1.429183e+02 | 22.659614 | 20.640489 | 1.714189 | 33.764922 |
| **TNFRI** | 2.462602e+02 | 2.795475e+03 | 268.892488 | 1.370583e+03 | 489.228828 | 1.973620e+03 | 561.853254 | 268.539201 | 407.255372 | 1229.287384 |
| **VEGFA** | 5.262186e+01 | 4.771512e+02 | 1055.665037 | 1.292433e+04 | 503.694935 | 5.652483e+03 | 278.677468 | 477.026744 | 377.963802 | 896.985279 |

Figure S7: Number of SASP proteins produced per cell per hour, from data in (Schafer et al., 2020)

**a**

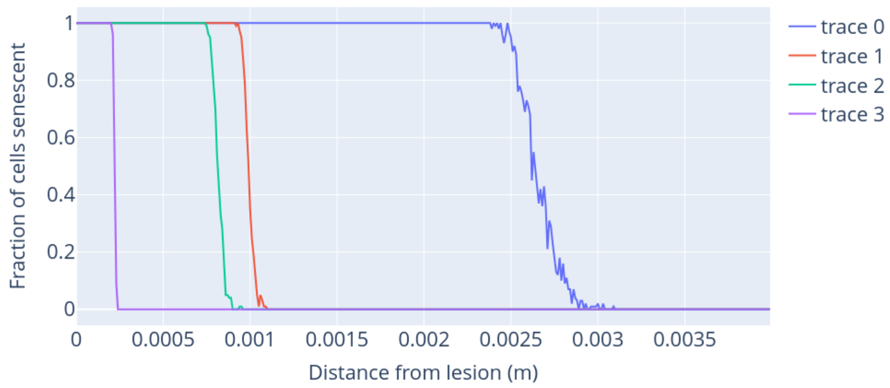


| Trace | Density | Binding rate ($k_{on}$ $M^{-1}$ $min^{-1}$) | Total number of receptors ($R_{tot}$) |
|---|---|---|---|
| Trace 0 | 0.35 | $1 \times 10^8$ | $10^4$ |
| Trace 1 | 0.35 | $1 \times 10^8$ | $10^5$ |
| Trace 2 | 0.2 | $1 \times 10^8$ | $10^5$ |
| Trace 3 | 0.35 | $5 \times 10^8$ | $10^5$ |

**b**

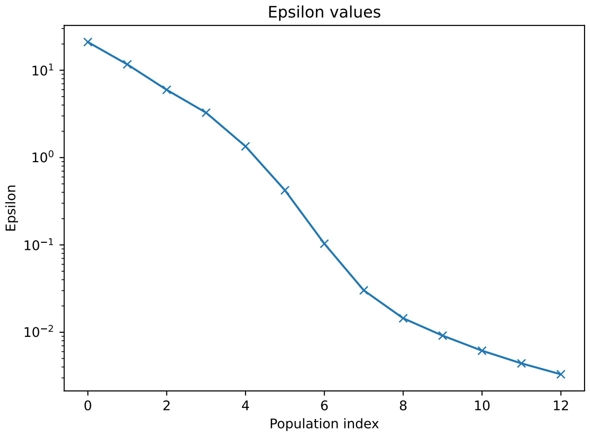


Figure S8: a) Posterior for the 4 inferred parameters, with red lines showing the true parameter values. True parameters were $r_{\text{cell}} = 10 \times 10^{-6}$, density = 0.35, length of experiment = 2 days, the radius of seeded region = $1000 \times r_{\text{cell}}$, $N_{\text{E}}$ = 2887 per hour, and $N_{\text{D}}$ = 140 per hour.
